# The Antiviral Enzyme, Viperin, Activates Protein Ubiquitination by the E3 Ubiquitin Ligase, TRAF6

**DOI:** 10.1101/2021.02.21.432004

**Authors:** Ayesha M. Patel, E. Neil G. Marsh

**Affiliations:** Department of Chemistry, University of Michigan, Ann Arbor, MI-48109; Department of Biological Chemistry, University of Michigan, Ann Arbor, MI-48109

## Abstract

Viperin is a broadly conserved radical SAM enzyme that synthesizes the antiviral nucleotide ddhCTP. In higher animals viperin expression also accelerates the degradation of various cellular and viral proteins necessary for viral replication; however, the details of this process remain largely unknown. Here we show that viperin activates a component of the protein ubiquitination machinery, which plays an important role in both protein degradation and immune signaling pathways. We demonstrate that viperin binds the E3 ubiquitin ligase, TRAF6, which catalyzes K63-linked ubiquitination associated with immune signaling pathways. Viperin activates ubiquitin transfer by TRAF6 ~2.5-fold and causes a significant increase in polyubiquitinated forms of TRAF6 that are important for mediating signal transduction. Our observations both imply a role for viperin as an agonist of immune signaling and suggest that viperin may activate other K48-linked E3-ligases involved in targeting proteins for proteasomal degradation.

## INTRODUCTION

Viperin (Virus Inhibitory Protein, Endoplasmic Reticulum-associated, Interferon iNducible) also denoted as cig5 and RSAD2 in humans,^1^ is strongly induced by type I interferons as part of the innate immune response to viral infection.^2–4^ Viperin is a member of the radical SAM enzyme superfamily and appears to be conserved in all 6 kingdoms of life,^5–7^ hinting at its ancient and ubiquitous role in combatting viral infection. Notably, viperin is one of the very few radical SAM enzymes found in higher animals.^8^ Viperin catalyzes the dehydration of CTP to form the antiviral nucleotide 3’-deoxy-3’,4’-didehydro-CTP (ddhCTP; Figure 1)^9^ through a radical mechanism initiated by reductive cleavage of SAM.^7, 10^ The antiviral properties of this nucleotide against RNA viruses derive from its ability to act as a chain-terminating inhibitor of some, but not all, viral RNA-dependent RNA polymerases.^9^

**FIGURE 1.**
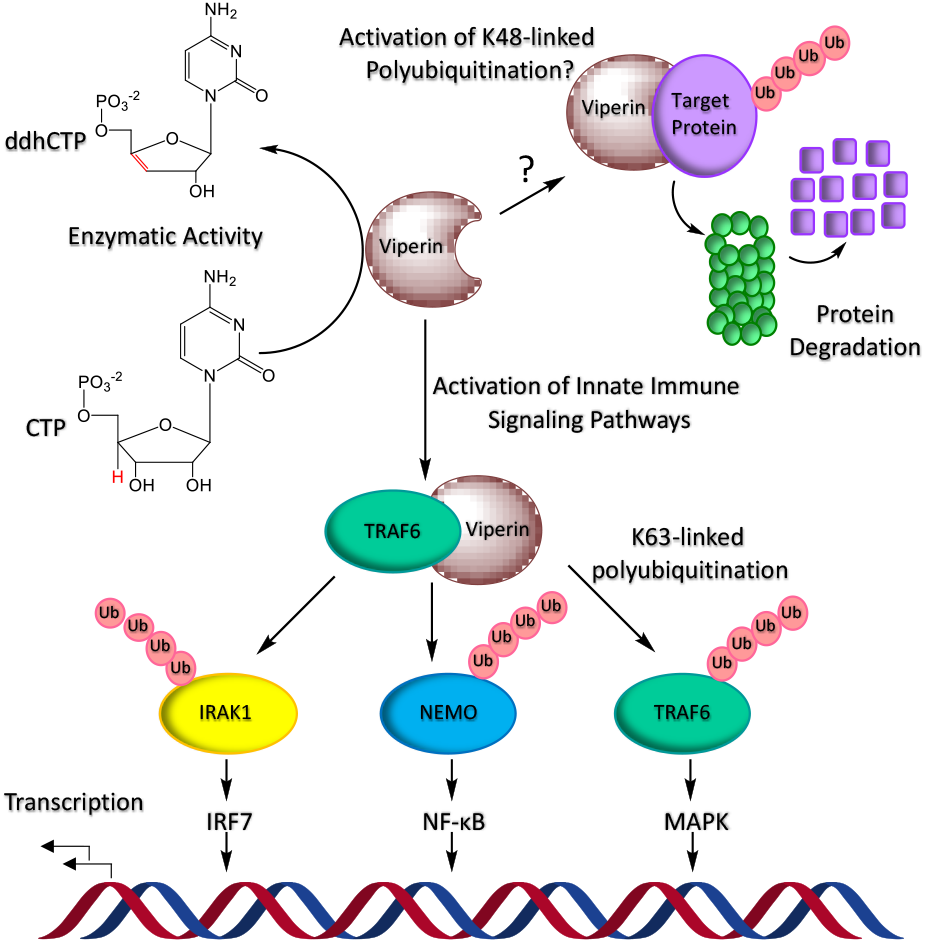
Overview of viperin’s interactions with the protein ubiquitination machinery and the E3-ligase,TRAF6.

In addition to synthesizing ddhCTP, viperin interacts with a wide range of cellular and viral proteins.^11–16^ This extensive network of protein-protein interactions remains poorly understood, but constitutes an equally important aspect of the enzyme’s antiviral properties. In many cases, it appears that viperin exerts its antiviral effects by facilitating the degradation of cellular and viral proteins important for viral replication.^11^ A prevailing view is that viperin recruits the protein ubiquitination machinery to target proteins for proteasomal degradation (Figure 1).^11, 17^ However, the evidence for viperin promoting protein ubiquitination is indirect and is based largely on studies using proteins transfected in mammalian cell lines.^11, 17^ Thus, it remains uncertain which components of the ubiquitination system viperin interacts with, or whether other proteins may be required for viperin to engage the ubiquitination system. Therefore to better understand viperin’s role in promoting protein ubiquitination, we have examined viperin’s interaction with the E3 ubiquitin ligase, TRAF6.^18–19^

The ubiquitination machinery comprises 3 enzymes:^20^ E1 is responsible for the ATP-dependent activation of ubiquitin and transferring it as its C-terminal thioester to various ubiquitin conjugating enzymes (E2). E2 enzymes interact with a large set of E3 ubiquitin ligases – ~700 in humans – that recognize different protein targets for ubiquitination.^21^

Polyubiquitin chains may be constructed through isopeptide bonds to various ubiquitin lysine residues. K48-linked polyubiquitination marks proteins for degradation by the 26S proteasome,^20, 22^ whereas Lys-63-linked polyubiquitination is important in activating various components of signal transduction pathways that trigger the immune response.^23^ E3 ligases, in particular, are highly regulated and may be activated or inhibited by a wide range of post-translational modifications and interactions with other proteins.^24^

TRAF6 (Tumor necrosis factor receptor-associated factor 6) is a member of the RING domain-containing E3 ligases.^25^ TRAF6 functions with the heterodimeric Ubc13/Uev1A E2-conjugating enzyme to synthesize K63-linked ubiquitin chains. TRAF6-mediated protein ubiquitination is central to several important signal transduction pathways,^22, 26^ including activation of the NF-κB pathway and the MAPK signaling cascade. These pathways regulate such diverse biological processes as cell growth, oncogenesis and immune and inflammatory responses.^26^ TRAF6 substrates include interleukin-1 receptor-associated kinases (IRAKs) and NF-κB essential modulator (NEMO).^27–28^ But TRAF6 also undergoes auto-ubiquitination specifically on Lys124^29^ and these polyubiquitinated forms serve to recruit and assemble downstream kinases and associated factors into signaling complexes that ultimately activate NF-κB and MAPK pathways.^30 26^

Recently, viperin was shown *in cellullo* to interact with TRAF6 to promote K63-linked polyubiquitination of interleukin receptor-associated kinase 1 (IRAK1) as part of innate immune signaling in the Toll-like receptor-7 and 9 (TLR-7/9) pathways.^15, 28^ The viperin-TRAF6 interaction provides a unique opportunity to test whether viperin functions as an activator of protein ubiquitination in a well-defined biochemical system. We have reconstituted the TRAF6 auto-ubiquitination system *in vitro* using purified enzymes. This has allowed us to demonstrate that viperin does indeed activate the E3 ligase activity of TRAF6, leading to a significant increase in the amount of polyubiquitinated TRAF6 species formed.

## RESULTS AND DISCUSSION

Full-length TRAF6 is a multidomain protein that forms large oligomers in the cell and has proven refractory to expression in *E. coli*. Therefore, to reconstitute the ubiquitination system *in vitro* we used a truncated TRAF6 construct comprising the RING and first 3 zinc-finger domains^19^ (designated TRAF6-N), which was previously shown to be functional and can be expressed and purified from *E. coli*.^19^ A human viperin construct lacking the first 50 residues of the ER-localizing N-terminal amphipathic helix, designated viperin-ΔN50, was expressed and purified from *E. coli* and the [4Fe-4S] cluster reconstituted as described previously.^16^ Preliminary pull-down experiments established that the truncated viperin-ΔN50 and TRAF6-N proteins form a stable complex with each other (Figure S1).

TRAF6 functions with the heterodimeric E2 ubiquitin-conjugating enzyme, Ubc13/Uev1A; this enzyme was expressed and purified from *E. coli* as described previously.^31^ The complete ubiquitination system was then reconstituted using commercially obtained E1 and ubiquitin. A typical assay comprised 0.1 μM E1, 2 μM Ubc13, 2 μM Uev1A, 2 μM TRAF6-N, and 2 μM viper-in-ΔN50 in 20 mM Tris-HCl buffer pH 7.5, 150 mM NaCl, 2 mM DTT, 2 mM ATP, and 5 mM MgCl_2_. Reactions were initiated by the addition of ubiquitin, 35 μM, and incubated at 37°C. At various times aliquots were removed and quenched by addition of SDS-PAGE loading buffer; samples were then analyzed by SDS-PAGE.

Initially we examined the activity of TRAF6-N in the absence of viperin, with a typical experiment shown in Figure 2. Under these conditions, the formation of diubiquitin was clearly visible after 4 min and tri-ubiquitin visible as a faint band after 8 min. At longer times, the formation of polyubiquitin is evident as a faint smear of higher molecular weight material. Control experiments established that in the background rate of ubiquitin ligation is negligible in the absence of TRAF6-N.

**FIGURE 2.**
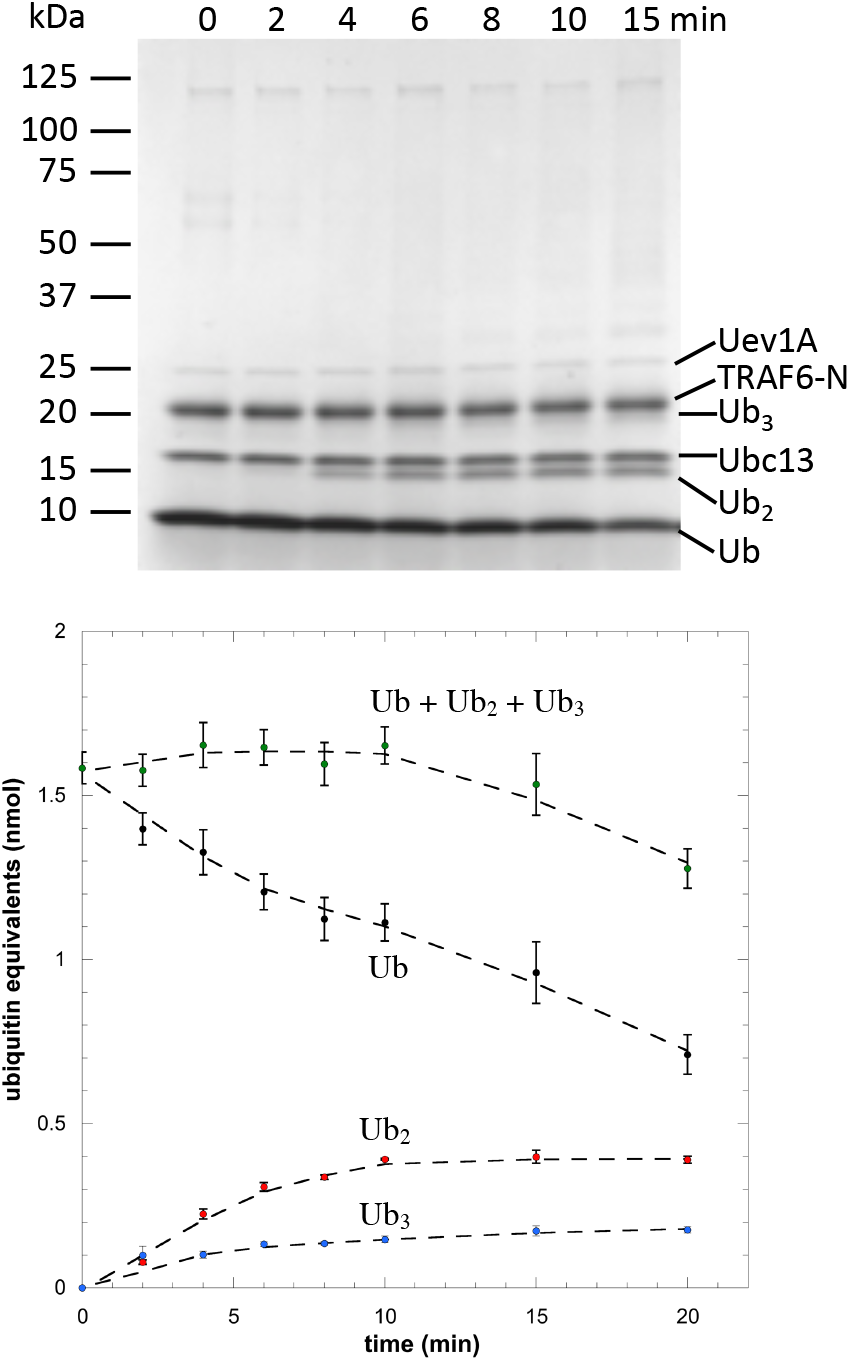
Kinetics of ubiquitin ligation catalyzed by TRAF6-N. *Top*: Representative Coomassie-stained gel showing consumption of ubiquitin and formation of ubiquitin oligomers. *Bottom*: Quantification of mono-, di- and tri-ubiquitin; after 20 min only a small fraction of the ubiquitin is converted to larger oligomers.

Quantification of the bands due to mono-, di- and triubiquitin by imaging of Coomassie stained gels (Figure 2) allowed the consumption of ubiquitin ligation to be quantified and the amount of ubiquitin incorporated into high molecular weight oligomers to be estimated. For the first 10 min of the reaction, the concentrations of di- and tri-ubiquitin increased and then plateaued. In contrast, the concentration of mono-ubiquitin steadily decreased as more ubiquitin was incorporated into high molecular weight oligomers. After 20 min ~ 80 % of the ubiquitin was accounted for by mono-, di- and triubiquitin, with only ~ 20 % converted to high molecular weight oligomers.

We then repeated the reaction with the addition of viperin-ΔN50 in a 1:1 ratio with TRAF6-N (2 μM of each enzyme). The addition of viperin markedly altered the kinetics of ubiquitination (Figure 3). In this case, there was an initial rapid increase in the amount of di-ubiquitin formed, which then decayed to a steady state level. Notably, ubiquitin was converted to high molecular weight species much more rapidly when viperin was bound to TRAF6-N. High molecular weight ubiquitin oligomers accounted for ~ 70 %, of the ubiquitin pool while mono-, di- and tri-ubiquitin comprised only ~ 30 %.

**FIGURE 3.**
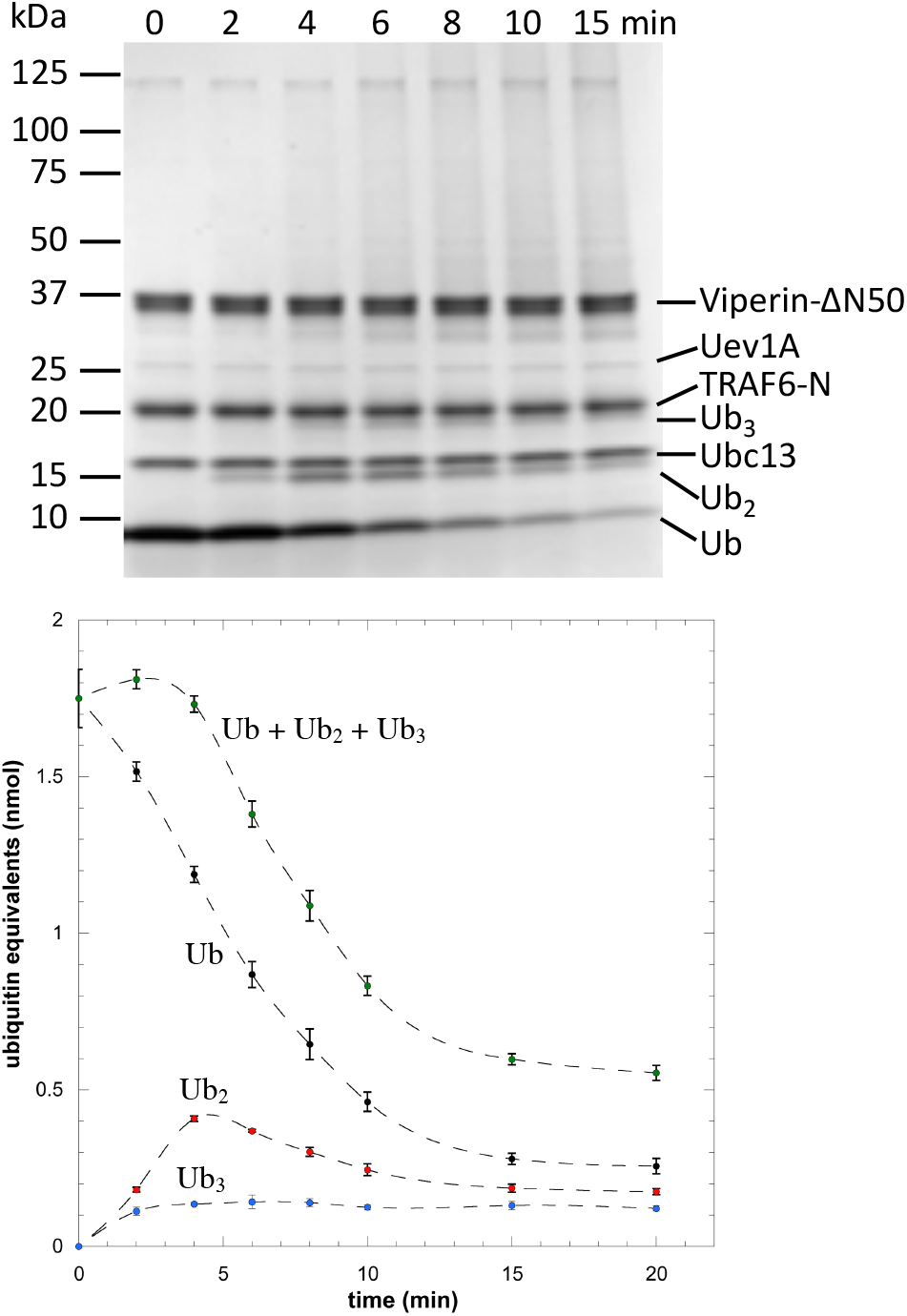
Activation of TRAF6-N by viperin. *Top*: Representative Coomassie-stained gel showing consumption of ubiquitin and formation of ubiquitin oligomers, note the smear of high M_r_ species at longer times. *Bottom*: Quantification of mono-, di- and tri-ubiquitin; these oligomers are rapidly depleted as they are converted to higher M_r_ species.

The iron-sulfur cluster of viperin appears to be important for TRAF6 activation, as viperin preparations in which the cluster had not been reconstituted did not activate TRAF6-N. Control experiments established that TRAF6-N activation is specific to viperin, as proteins such as bovine serum albumin had no effect on TRAF6-N activity. These results clearly demonstrate that viperin activates TRAF6-N and promotes the formation of longer polyubiquitin chains that are considered to be important mediators of signaling in the MAPK and NF-kB pathways.^26, 30^

TRAF6 is known to auto-ubiquitinate Lys124,^29^ a process that is important for its role in signal transduction.^30^ To examine whether viperin promotes TRAF6 auto-ubiquitination, we probed gels with antibodies against ubiquitin and the N-terminal domain of TRAF6 (Figure 4). Immunoblotting with anti-ubiquitin antibody confirmed identity of the di- and tri-ubiquitin bands and, as expected, strongly stained the high molecular weight material evident in Coomassie-stained gels. The high molecular weight material also cross-reacted with anti-TRAF6 antibodies demonstrating it represents auto-ubiquitinated forms of TRAF6-N. When probed with anti-viperin antibodies, only the viperin band was crossreactive. This result demonstrates that although viperin promotes TRAF6 polyubiquitination, it is not itself a substrate for ubiquitination.

**FIGURE 4.**
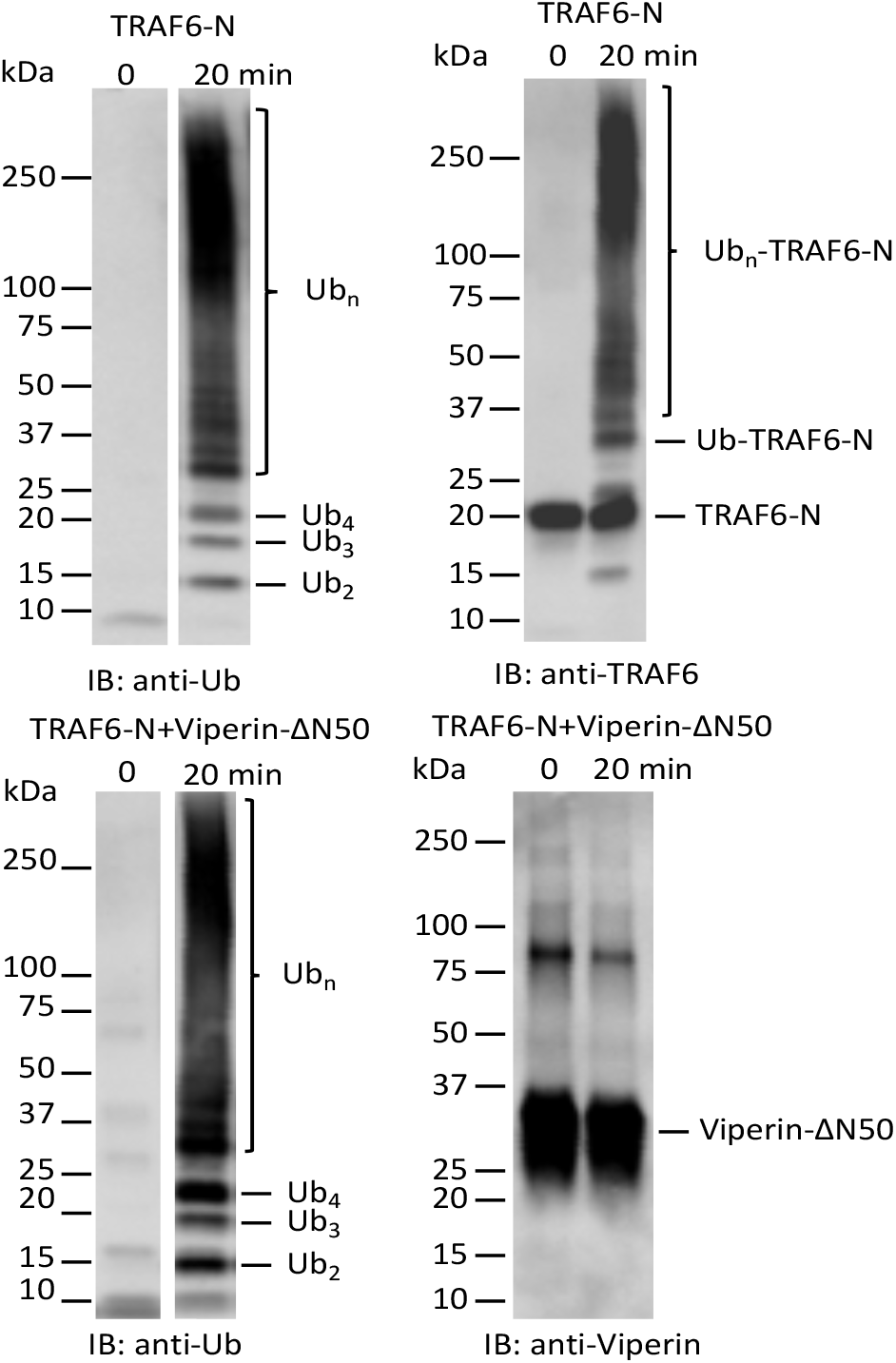
Immunoblot analysis of ubiquitination reactions. *Top* Staining for ubiquitin (left) and TRAF6 (right) in reactions containing TRAF6-N. *Bottom:* Staining for ubiquitin (left) and viperin (right) in reactions containing TRAF6-N and viperin.

Although the time course for ubiquitin ligation is complex, at early time points the major reaction catalyzed by TRAF6-N is the formation of di-ubiquitin:

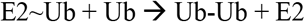

The ubiquitin-charged E2 functions as a substrate for TRAF6-N, which is then rapidly replenished through the action of E1 so that the steady state concentration of E2~Ub remains constant. This simplification allowed us to quantify the ubiquitin ligase activity of TRAF6-N and compare its activity when complexed with viperin. Preliminary experiments established that the rate of ubiquitin consumption was linear with TRAF6-N concentration (Figure S2). Under these conditions, the apparent turnover number for di-ubiquitin formation by TRAF6-N was *k*_app_ = 0.47±0.06 min^−1^, whereas in the presence of viperin *k*_app_ = 1.25±0.08 min^−1^ representing a ~ 2.5-fold rate enhancement (Figure S3).

TRAF6 is one of the better studied members of this class E3 ligases, in part due to the important role it plays in NF-kB and MAPK signaling.^22, 26^ However, to our knowledge, quantitative measurements of rate at which TRAF6 catalyzes ubiquitin transfer have not been previously been reported. The kinetics of ubiquitin transfer catalyzed by various other E3 ligases have been quite extensively investigated, with kcat ranging from several per second, e.g. the RING-E3 ligase SCF^Cdc4^ ^32^ and HECT-E3 ligase E6AP^33^, to several per minute, e.g. the RING-E3 ligase San1.^34^ Compared with these E3 ligases, the rate of ubiquitin ligation catalyzed by TRAF6-N is relatively slow, but we note that ubiquitination rates are also dependent on the protein substrate and may accelerate as the polyubiquitin chain is extended.^32^ Furthermore, TRAF6-N lacks the C-terminal TRAF domain through which TRAF6 binds many of its protein substrates and which may also influence the ligase activity of the enzyme.

A role for viperin in immune signaling was initially suggested through studies on TRAF6-catalyzed polyubiquitination of IRAK1 in mouse cell-lines lacking viperin.^28^ More recently, our studies in HEK 293T cells demonstrated that co-transfection of viperin with TRAF6 significantly increased the polyubiquitination of IRAK1.^15^ However, these studies left open the possibility that viperin activated TRAF6 indirectly through additional unknown factor(s). Reconstituting the ubiquitination system *in vitro* with purified enzymes has allowed us to unambiguously demonstrate viperin’s role in activating TRAF6. Viperin both speeds up the rate of ubiquitin consumption and increases the formation of high molecular weight auto-ubiquitinated forms of TRAF6 that mediate downstream signaling. Although the ~2.5-fold activation of TRAF6 by viperin is relatively modest, this level of amplification may be appropriate to modulating transcription of the various genes needed to establish the antiviral response.

Here we have shown that viperin interacts with the N-terminal RING-domain of TRAF6, whereas it is known that the C-terminal TRAF domain (lacking in TRAF6-N) mediates TRAF6’s interactions with most other protein substrates.^24^ This modular arrangement suggests that activation of TRAF6 E3-ligase activity by viperin may enhance polyubiquitination of other target proteins, which would be consistent with our observation that viperin stimulates TRAF6-catalyzed polyubiquitination of IRAK1.^15^ These observations provide further support the idea, for which there is extensive but indirect evidence in the literature, that viperin, more broadly, activates other K48-linked E3 ligases to increase proteasomal degradation of specific proteins in response to viral infection.

## EXPERIMENTAL METHODS

### Plasmids, reagents, and antibodies

Human TRAF6-N (aa 50-211) construct in pET21c was a kind gift of Prof. Hao Wu (Harvard University). Ubc13 and Uev1A constructs in pGEX6P3 were a kind gift of Prof. Catherine Day (University of Otago, New Zealand). The truncated human viperin-ΔN50 (aa 51-361) was cloned into pRSF-duet vector with an N-terminal His tag. The human E1 enzyme (E-304-050) was purchased from R&D systems. Ubiquitin was purchased from Sigma Aldrich. Pierce protein A/G plus agarose resin and control agarose resin (Pierce classic IP kit 26146) were purchased from Thermo-Fisher Scientific. The rabbit polyclonal viperin antibody (11833-1-AP), rabbit polyclonal ubiquitin antibody (10201-2-AP) both were obtained from ProteinTech. The rabbit polyclonal TRAF6 antibody (sc-7221) was obtained from Santa Cruz Biotechnology. Goat anti-rabbit (170-6515)) Ig secondary antibody was purchased from BioRad.

### Protein Expression and Purification

#### TRAF6-N

Human TRAF6-N^18^ containing a C-terminal His tag was expressed in *E. coli* BL21 (DE3). Cultures were grown to OD_600_ 0.6-0.8 before cold shocking the cells in ice bath for 30 min. The protein was induced with 0.5 mM IPTG and 0.1 mM ZnCl_2_ and incubated at 20°C overnight. Cells were lysed by sonication in buffer A (50 mM Tris-HCl pH 8.0, 300 mM NaCl, 10 mM imidazole and 10% glycerol) with protease inhibitor cocktail and clarified by centrifugation at 18,000 rpm for 1 h at 4°C. The cleared lysate was loaded onto the HisTrap prepacked column. The column was washed with buffer A (10 mM imidazole) until no further protein eleuted. The column was then washed with 90% buffer A and 10% buffer B (50 mM Tris-HCl pH 8.0, 300 mM NaCl, 500 mM imidazole and 10% glycerol) followed by washing with 80% buffer A and 20% buffer B. Finally, the protein was eluted with 60% buffer A and 40% buffer B (200 mM imidazole). The fractions containing TRAF6-N were pooled, concentrated, and further purified by size-exclusion chromatography. The concentrated protein was loaded on to Superdex 200 16/120 column equilibrated in 50 mM Tris-HCl pH 8.0, 150 mM NaCl, and 10% glycerol. Fractions containing TRAF6-N were pooled, concentrated and stored at – 20 C.

#### Ubc13 and Uev1A

The expression, purification, and GST tag cleavage protocols of Ubc13 and Uev1A were performed as described previously.^35^

#### Viperin-ΔN50

Human viperin-ΔN50 was cloned into the expression vector pRSF-duet, and transformed into *E. coli* BL21 (DE3) using standard methods. The expression of viperin-ΔN50, purification of the protein and reconstitution of the iron-sulfur cluster under anaerobic conditions were performed as described previously.^12^

#### Co-immunoprecipitation of viperin and TRAF6-N

In an anaerobic environment (Coy anaerobic chamber), 1 μM viperin-ΔN50 and or 1 μM TRAF6-N were mixed in a buffer containing 50 mM Tris-HCl pH 8.0, 150 mM NaCl, 5 mM DTT, and 10% glycerol and incubated at 4 °C for 1 h. Then, 0.5 μg of anti-viperin antibody was added to each reaction and incubated at 4 °C for further 1 h. Next, 10 uL of protein A/G Agarose beads (20 uL slurry) equilibrated in the same buffer, were added and the mixture was then incubated with end-to-end mixing outside the Coy chamber in 4 °C for 1 h. The beads were washed three times with excess buffer before incubating with 2X SDS loading buffer containing 5% 2-mercaptoethanol. The mixture was then agitated for 20 min, and heated at 95 °C for 10 min, and the beads removed by centrifugation. The proteins were analyzed by SDS-PAGE (4 – 20 % gradient gels) and immunoblotted with appropriate antibodies using standard protocols.

### Ubiquitination assay

All assays containing viperin-ΔN50 were conducted inside a Coy anaerobic chamber. For ubiquitination assays, 0.1 μM E1, 2 μM Ubc13, 2 μM Uev1A, 2 μM TRAF6-N, and/or 2 μM viperin-ΔN50 were mixed in a buffer containing 20 mM Tris-HCl pH 7.5, 150 mM NaCl, 2 mM DTT, 2 mM ATP, and 5 mM MgCl_2_ and incubated at 37°C. Reactions were initiated by the addition of ubiquitin, 35 μM final concentration, and at various times, aliquots of the assay mixture were removed and quenched by adding equal volumes of 2X SDS loading buffer containing 2-mercaptoethanol. Proteins were analyzed by SDS-PAGE on 4-20% gels. Gels were stained with Coomassie brilliant blue and ubiquitin bands quantified with reference to known standards.

To examine which bands represented ubiquitinated forms of TRAF6, samples were analyzed using by SDS-PAGE on 4-20% gels that were then subjected to immunoblot analysis using standard techniques with antibodies to ubiquitin, viperin or the N-terminal domain of TRAF6.

## AUTHOR INFORMATION

### Author Contributions

E.N.G.M. conceived the project; A.M.P and E.N.G.M. designed the experiments; A.M.P. performed the experiments; A.M.P and E.N.G.M. wrote the paper.

## ACKNOWLEDGMENT

We thank Prof. Hao Wu (Harvard University) and Prof. Catherine Day (University of Otago) for their kind gifts of expression vectors for TRAF6-N and Ubc13/Uev1A respectively.

